# Systematic Pan-cancer Functional Inference and Validation of Hyper, Hypo and Neomorphic Mutations

**DOI:** 10.1101/2023.04.29.538640

**Authors:** Somnath Tagore, Samuel Tsang, Gordon B. Mills, Andrea Califano

## Abstract

While the functional effects of many recurrent cancer mutations have been characterized, the TCGA repository comprises more than 10M non-recurrent events, whose function is unknown. We propose that the context specific activity of transcription factor (TF) proteins—as measured by expression of their transcriptional targets—provides a sensitive and accurate reporter assay to assess the functional role of oncoprotein mutations. Analysis of differentially active TFs in samples harboring mutations of unknown significance—compared to established gain (GOF/hypermorph) or loss (LOF/hypomorph) of function—helped functionally characterize 577,866 individual mutational events across TCGA cohorts, including identification of mutations that are either neomorphic (gain of novel function) or phenocopy other mutations (*mutational mimicry*). Validation using mutation knock-in assays confirmed 15 out of 15 predicted gain and loss of function mutations and 15 of 20 predicted neomorphic mutations. This could help determine targeted therapy in patients with mutations of unknown significance in established oncoproteins.

## Introduction

Tumorigenesis is characterized by the progressive accrual of mutations in oncogenes and tumor suppressors, leading to transformation events^1^. Remarkably, however, the vast majority of samples in large scale repositories still lacks a genetic rationale for tumorigenesis, due either to the non-recurrent (*i.e., private*) nature of their mutations or to presence of multiple mutations, each one providing only a modest contribution to tumorigenesis (*i.e., field effect*). Indeed, recent studies have shown that the additive effect of a large number of non-recurrent, or poorly characterized mutational events, with weak functional effects on an individual basis, can induce massive dysregulation of cell state followed by transformation^2^. As a result, differentiating neutral mutations, also known as passenger mutations, from mutations that may play a relevant, albeit small or uncharacterized functional effect, remains a highly relevant and still largely elusive question. In addition, with an increasing repertoire of small molecular inhibitors targeting oncogenic proteins and their downstream mediators, the question of whether a patient with a mutation of unknown significance in an established oncogene should be treated with the corresponding inhibitor is highly translationally relevant.

Gene mutations can either be pathogenic or have no deleterious effects on the disease of interest, as mediated by the protein they encode. In terms of functional mutations, gain (GOF) and loss of function (LOF) events—also called *hypermorph* and *hypomorph* mutations—can increase or abrogate the normal physiologic activity of a protein. When GOF or LOF events occur in oncogenes or tumor suppressors, respectively, they can be pathogenic. A third class of neomorphic events (NEO) has also been reported^3^, however. These mediate the emergence of entirely novel functions, unrelated to the physiologic function of the wild type (WT) protein, for instance by implementing novel protein-protein interactions that disrupt normal cell behavior.

An inventory of genetically profiled tumor sample repositories reveals more than ten million non-recurrent, yet potentially actionable genetic alterations in established oncoproteins and tumor suppressors, whose functional effect are yet to be established^4^. Functional characterization of these events is critically important, not only to improve our molecular level understanding of cancer but also from a translational perspective, since it can help assess whether patients harboring these mutations would benefit from available targeted inhibitors or whether their use may be ineffective or even harmful. For instance, targeting a hypomorph or neomorph mutation in PIK3CA with a targeted inhibitor may be ineffective, or even harmful if it induced aberrant activation of alternative oncogenic pathways^5^. Similarly, pharmacologic targeting of neomorphic mutations with an inhibitor of the WT protein’s function may fail to reduce or even increase their tumorigenic effect^6^. The challenge of functionally characterizing mutations of unknown significance extends to (a) mutations that may have been functionally characterized in one tumor context but are not well studied in another context, where they may have substantially different functional impact^7^, (b) co-mutations that may affect the functional role of the primary mutated oncogene^7^, or (c) mutations that may phenocopy the functional effects of established oncogenes (*mutational mimicry*)^8^.

While there are highly effective experimental tools for the functional characterization of cancer mutations^7, 53^, scaling these methodologies to cover the full repertoire of somatic mutations—for instance, using saturation mutagenesis assays^9^—is exceedingly laborious and expensive. In addition, since the repertoire of rare/private mutations that may be observed in future patients cannot be predicted ahead of time, the relatively slow nature of available approaches for experimental assessment of mutation function and pathogenicity would preclude real-time patient management. This is especially relevant because tumor context plays a critical role in determining the functional relevance of specific mutations, thus requiring that each mutation be studied in the specific cellular context in which its function should be assessed^10^.

Most transforming mutations occur either in signal transduction pathways or in transcriptional regulators^11^. We have shown that aberrant, mutation-induced behavior of proteins in both of these classes can be effectively characterized in terms of its either physical or indirect effects on transcription factor (TF) and co-factor (co-TF) activity, which either represent the ultimate downstream effectors of aberrant signaling^12^ or are directly affected by mutations, including in their non-coding regulatory regions^13^. We have also shown that differential activity of TF and co-TF proteins can be accurately measured by leveraging their transcriptional targets as a multiplexed gene reporter assay, using the VIPER algorithm^14^. As a result, we hypothesize that hyper, hypo, and neo-morphic mutations will induce highly specific TF/co-TF activity signatures that can be leveraged to characterize mutations of unknown significance.

Based on this assumption, we developed PHNToM (**P**rotein-activity based identification of **H**ypermorph, Hypomorph, **a**nd **N**eomorph, **T**herapeutically-relevant **M**utations), a machine learning algorithm that first identifies TF activity signatures induced by GOF and/or LOF mutations and then uses them to accurately classify mutations of unknown significance in the same gene, as either neutral, hypermorphic, hypomorphic, or neomorphic, as well as to characterize their ability to mimic the effect of mutations in other genes (*mutational mimicry*).

Of the 10 million events of unknown significant in TCGA, PHNToM analysis helped functionally characterize 577,866 mutational events in 25 cohorts represented in The Cancer Genome Atlas (TCGA)^15^. Of these, 282,100, 191,230, 102,123 and 2,413 were predicted to represent hyper, hypo, neutral and neomorphic events, respectively, thus greatly increasing the repertoire of mutations with functional assessment. The remaining events occurred either in genes that had too few previously characterized GOF/LOF mutations and could thus not be used to “learn” a TF/co-TF activity signature or in cohort too small to support the analysis.

To validate prediction accuracy, we ectopically expressed a subset of mutated transgenes in wild-type (WT) cells and measured their effect on protein activity based on analysis of PLATE-Seq^16^ profiles. These assays confirmed the high accuracy and sensitivity of the methodology, including validation of 15 out 15 (100%) predicted GOF and LOF mutations and 15 of 20 (75%) predicted neomorphic mutations. Based on the confirmed accuracy of the predictions, we also produced extensive characterization of candidate mutational mimicry events. In addition, by associating mutations in each oncoprotein to a specific TF/co-TF signature, the analysis provides novel insight into the tumor-specific pathways affected by its pathologic function.

We first selected *PIK3CA* and *TP53* mutations in the TCGA breast cancer cohort (BRCA) as an illustrative example of the approach. Both of these are well-studied genes with extensive functional characterization. As reported in cBioPortal^17, 18^, the BRCA cohort comprises 1,084 samples, of which 333 and 352 harbor *PIK3CA* and TP53 mutations, respectively. Yet, of these, only 124 and 154 have been characterized in terms of their functional significance, respectively. We then report the analysis for the full repertoire of mutations that could be analyzed by the proposed methodology.

## Results

PHNToM was designed to systematically predict the effect of mutations of unknown significance in established oncoproteins and tumor suppressors (oncoproteins for short), in a specific tumor context. To accomplish this goal, it leverages VIPER^14^, an extensively validated algorithm designed to quantitatively assess the transcriptional activity of ∼3,700 regulatory proteins^2, 19, 20^— comprising transcription factor (TF) and co-factor (co-TF) proteins (TF for short), as defined in Gene Ontology^21^—based on the differential expression of their tissue-specific transcriptional targets, as identified by the equally extensively validated ARACNe algorithm^22–24^. The rationale for this approach is that established GOF or LOF mutations will induce highly specific TF activity patterns that can be learned and then used to predict the effect of uncharacterized mutations in the same protein. A conceptual methodological workflow is provided in **Figure 1**.

**Figure 1:**
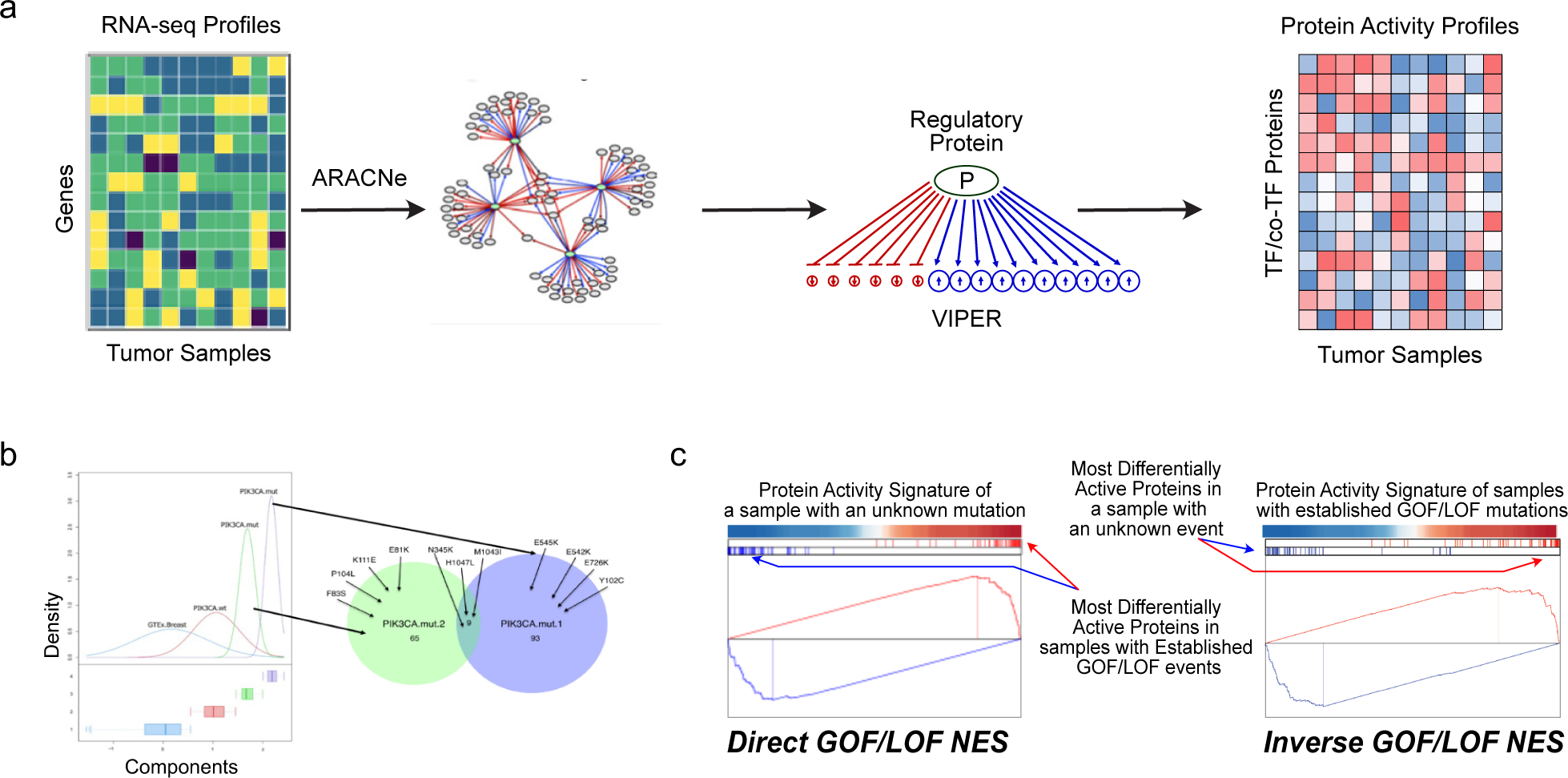
Conceptual overview of the PHNToM algorithm. (a) Generation of TF/co-TF protein activity profiles. A regulatory network—including positively regulated and repressed targets of all TF and co-TF proteins (regulon)—is first generated by ARACNe-AP analysis of RNA-seq profiles from a specific tumor cohort. Then, the VIPER algorithm is used to assess the activity of each TF and co-TF based on the expression of its regulon. **(b) Multi-modal Gaussian mixture model.** For each mutation-harboring gene under consideration, a multi-modal analysis of WT, mutated, and GTEx samples (either tissue matched if available or averaged over the entire GTEx repository) is performed to identify distinct modes associated with aberrant protein activity, as assessed by VIPER. For instance, the four curves represents the PIK3CA activity probability density in (*i*) GTEx normal breast samples (blue), (*ii*) *PIK3CA^W^*^T^ samples in the TCGA BRCA cohort (red), and (*iii*) two *PIK3CA^Mut^*sample subsets—as identified by multi-modal analysis of the BRCA cohort in TCGA—with either intermediate (green) or high (purple) PIK3CA activity, as also summarized by the Venn diagrams. Negative controls for the analysis are selected as the subset of WT samples whose PIK3CA activity is indistinguishable from GTEx samples (i.e., p > 0.05, based on Student’s T-test). **(c) Protein Activity Enrichment Analysis (PAEA).** Two enrichment analyses are performed: (i) the Direct GOF/LOF NES (DGN) represents the enrichment of the 50 most differentially active/inactive proteins (25 + 25) of a sample harboring a mutation of unknown significance (test sample) in proteins differentially active in the Consensus GOF/LOF Signature and (*ii*) the Inverse GOF/LOF NES (IGN) represents the enrichment of 50 most differentially active and inactive proteins in the Consensus GOF/LOF Signature, in proteins differentially active in the sample harboring an unknown significance mutation (test sample).

Briefly, given a protein of interest (P) in a specific tumor type, we generate an activity signature associated with its mutated state by comparing samples harboring the event in a tumor cohort (P^Mut^) to a subset of wild-type controls from the same cohort (P^WT^). In the following paragraphs, we will discuss how both P^Mut^ and P^WT^ samples are selected to achieve optimal results.

### Cohort Selection and Protein Activity Analysis

Since ARACNe requires ≥ 100 gene expression profiles to accurately infer protein regulons, our analysis was limited to 25 TCGA cohorts of appropriate size (**Suppl. Tables 1,2**). Once protein regulons are inferred, VIPER is used to transform any tissue specific gene expression profile into a *protein activity* profile^14^ (**Figure 1a, Suppl. Figure 1a**). A large body of literature has shown that VIPER-assessed protein activity is highly accurate and compares favorably with antibody-based measurements, even when using single cell profiles^14, 20, 25^.

### Negative Control Sample Selection

To generate appropriate controls representing the effect of the WT protein in a specific tumor-subtype, we start by selecting all tumor-specific P^WT^ samples. However, since protein activity can be dysregulated by overexpression, genomic amplification/loss or a variety of mutational events, or aberrant paracrine and endocrine signals in their upstream pathways—all frequent events in cancer—these samples may still present aberrant activity of the protein of interest, thus producing a significant confounding factor.

For instance, while *PIK3CA^Mut^* samples in breast cancer are generally associated with high VIPER-assessed activity of the encoded protein, *PIK3CA*^WT^ samples are also significantly biased towards higher PIK3CA activity, compared to normal epithelial breast samples in GTEx^26^ (**Figure 1b**). For this purpose, differential PIK3CA activity was computed by VIPER using the centroid of the entire GTEx repository as a reference. Thus, indiscriminate use of *PIK3CA*^WT^ samples in the TCGA BRCA cohort as negative controls would negatively affect the analysis.

To address this challenge, PHNToM restricts negative controls to a subset of *PIK3CA*^WT^ BRCA samples, whose PIK3CA activity is not statistically significantly different from those in the GTEx epithelial breast tissue cohort (p > 0.05, by Student’s T-test) (see STAR Methods: *P^WT^*^0^ *Null Model Generation*). We call these refined control samples the P^WT0^ set (i.e., *PIK3CA^WT^*^0^ in this case).

A second critical confounding factor is introduced by the non-random stratification of mutations across transcriptionally distinct tumor subtypes in the same tumor cohort. For instance, *PIK3CA*^Mut^ samples are significantly enriched in hormone receptor positive (HR+) and HER2+ samples compared to triple negative breast cancer (TNBC) samples. As a result, any signature obtained by comparing *PIK3CA*^Mut^ vs. *PIK3CA*^WT^ samples would be heavily biased by differential TF activity in the corresponding subtypes. While this may be mitigated by performing this analysis on an individual tumor subtype basis, for instance based on cluster analysis, there may be finer-grain stratification structures that may be missed by such an approach, thus producing significant confounding effects. Thus, rather than using all P^WT0^ samples as controls, PHNToM selects only those representing the k-nearest neighbors of the specific P^Mut^ sample being analyzed, using a protein activity distance metric, as implemented by the OncoMatch algorithm^19^ (see STAR Methods: *K-nearest neighbor analysis)*. For the analyses reported in this manuscript, *k* = 5 is used to ensure appropriate assessment of VIPER statistics, see^14^. For simplicity, we will call these negative control samples the P<u>^WT^</u> set.

### Generation of a Consensus GOF/LOF signature

For each protein with an established set of functionally characterized mutations, we generate a Consensus GOF/LOF Signature (*Consensus Signature*, for short) as a TF list ranked by the statistical significance of their VIPER-assessed differential activity in P^Mut^ vs. P<u>^WT^</u> samples, integrated across all samples harboring GOF or LOF mutations (see STAR Methods: *Consensus GOF/LOF Signature*). To integrate GOF and LOF into a single signature, the protein activity sign is inverted for LOF events. MR overlap between the Consensus Signature and equivalent signatures generated by comparing a sample harboring a mutations of unknown significance to the P<u>^WT^</u> samples, by OncoMatch analysis, is then used to assess its GOF, LOF, NEO, or neutral status, as discussed in the next section.

### Predicting Mutation Functional Significance

To classify mutations as GOF, LOF, neomorphic or neutral, PHNToM performs Protein Activity Enrichment Analysis (PAEA). This represents a straightforward extension of GSEA^27^ that replaces differential gene expression with differential, VIPER-assessed protein activity. Results are based on the joint analysis of two PAEA normalized enrichment scores (NES):

***Direct GOF/LOF NES (DGN)***: this NES represents the enrichment of the most differentially active proteins—top 25 and bottom 25—of a sample harboring a mutation of unknown significance (test sample) in proteins differentially active in the Consensus Signature (**Figure 1c**).

***Inverse GOF/LOF NES (IGN):*** this NES represents the enrichment of the most differentially active proteins in the Consensus Signature—top 25 and bottom 25—in proteins differentially active in a sample harboring a mutation of unknown significance (test sample) (**Figure 1c)**.

We use the top and bottom 25 most differentially active TFs in this analysis because we have shown that, on average, such TF repertoire is sufficient to canalize the effect of the vast majority of mutations in TCGA cohorts^2^. However, we have also shown that predictions remain virtually unaffected when this number ranges from 10 to 200, thus supporting the robustness of the analysis^28^. The following four scenarios are thus possible:

1. ***The mutation is predicted as a GOF event*** if both the DGN and IGN are positive and statistically significant (*p* ≤ 0.05, FDR corrected) and their difference, |DGN - IGN|, is not statistically significant.
2. ***The mutation is predicted as a LOF event*** if both the DGN and IGN are negative and statistically significant (*p* ≤ 0.05) and their difference, |DGN - IGN|, is not statistically significant.
3. ***The mutation is predicted as neomorphic (NEO) event*** if the difference between the DGN and IGN, s_NEO_ = |DGN - IGN|, is statistically significant. This may happen, for instance, if the DGN is statistically significant while the IGN is either non-significant or has the opposite sign. However, it may also happen if both metrics are significant and concordant, yet substantially different. The rationale for nominating neomorphic events based on this metric is that, by definition, neomorphic events will dysregulate many TFs that are not affected by established GOF and LOF events, thus resulting in inconsistent direct and indirect enrichments.
4. ***The event is a neutral (NEU) mutation*** if neither metric is statistically significant.

For GOF and LOF events, statistical significance p(GOF; LOF) is assessed by Stouffer’s integration of the DGS and IGS values. For NEO events, statistical significance, p(NEO), is assessed by the absolute value of the difference between DGN and IGN values, using a null model based on the probability density of the s_NEO_ metric across all established GOF and LOF events (gold standard dataset, see STAR Methods: *Mutations Selection*). Thus, each event (either established or of unknow significance) is associated with two p-values, one representative of its potential GOF/LOF nature p(GOF; LOF) and one associated with its potential neomorphic activity p(s_NEO_).

The four different outcomes are illustrated in **Figure 2a-d**. For instance, *PIK3CA^H1047R^* (TCGA.EW.A1PC) is predicted as a GOF event, since it shows significant positive DGN (NES = 15.2, p = 2.9×10^−52^) and IGN (NES = 10.2, p = 2.1×10^−24^) (**Figure 2a**), while *PIK3CA^D350N^* (TCGA.AC.A2FF) is predicted as an LOF event, because it shows significant negative DGS (NES = −10.8, p = 3.1×10^−27^) and IGN (NES = −7.84, p = 4.3×10^−15^) (**Figure 2b)**. In contrast, *PIK3CA^E545K^* (TCGA.E9.A1NG) is predicted as a NEO event because it shows significant positive DGS (NES = 3.36, p = 7.8×10^−4^) and IGN (NES = 0.68, p = 1.04×10^−1^) (**Figure 2c)**. Finally, *PIK3CA^E^*^103*_G106delins*^ (TCGA.C8.A26X) is predicted as a neutral event since its DGN and IGN are not significant (NES = 2.48, p = 1.3×10^−2^ and NES = 1.6, p = 1.2×10^−1^) (**Figure 2d)**.

**Figure 2:**
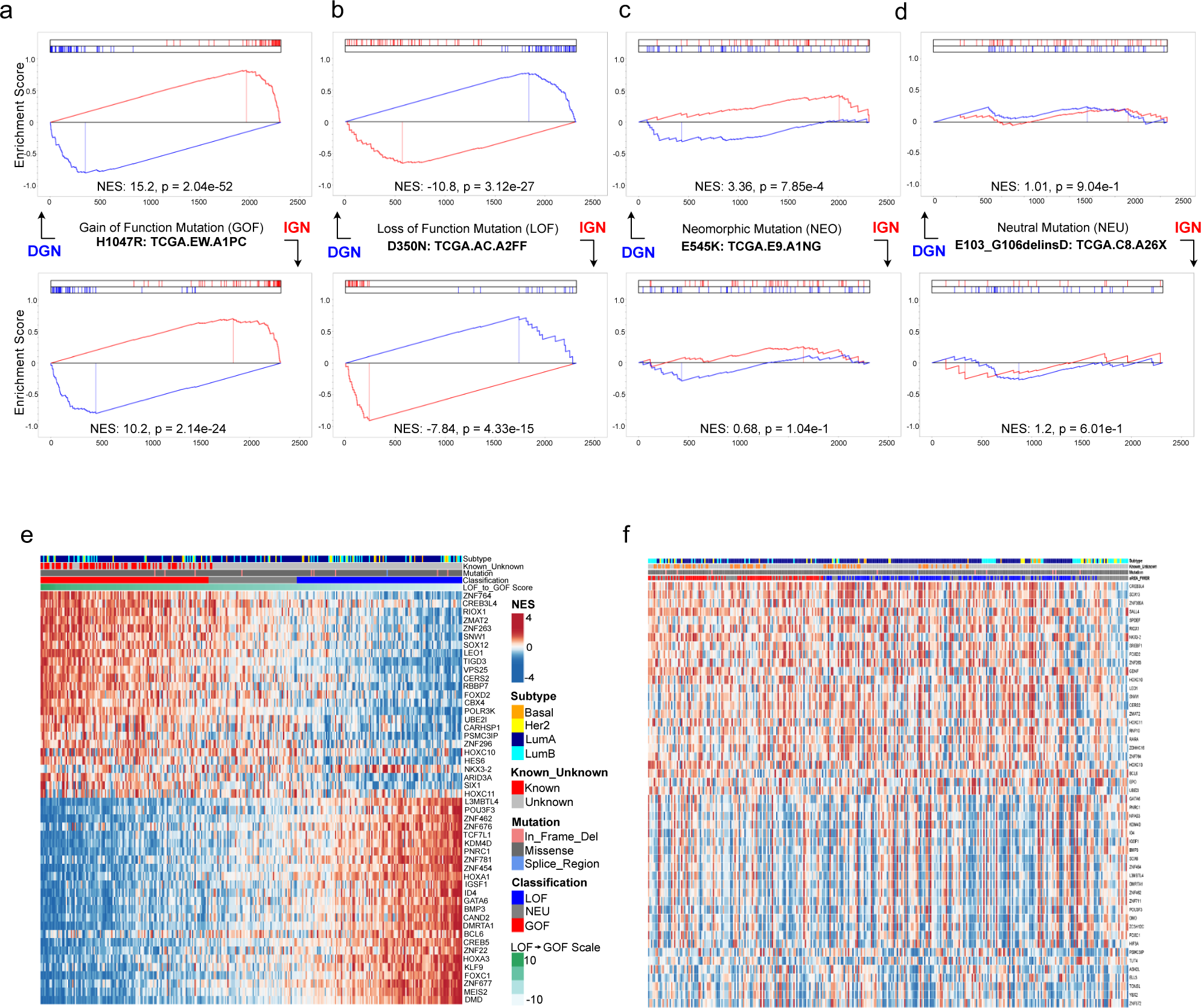
PHNToM analysis of PIK3CA mutations in the TCGA breast cancer cohort (BRCA) **(a-d)** DGN and IGN analysis of representative hypermorphic (GOF), hypomorphic (LOF), neomorphic (NEO) and neutral (NEU) PIK3CA mutations. The analysis shows significant positive DGN and IGN for *PIK3CA^H1047R_TCGA.EW.A1PC^* (GOF), significant negative DGN and IGN for *PIK3CA^D350N_TCGA.AC.A2FF^* (LOF), significant positive DGN but not IGN for *PIK3CA^E545K_TCGA.E9.A1NG^*(NEO), and not significant DGN and IGN for *PIK3CA^E^*^103*_G106delinsD_TCGA.C8.A26X*^ (NEU). **(e)** Heatmap showing *PIK3CA*^Mut^ BRCA samples (columns), sorted from the most hypermorphic to the most hypomorphic one based on PHNToM cross-validation analysis. The 25 most differentially active and 25 most differentially inactive TF/co-TF proteins in the consensus GOF/LOF signatures, representing candidate PIK3CA effectors, are shown in the rows. The Consensus GOF/LOF signature was generated by averaging the differential activity of TFs and co-TFs across all samples harboring established GOF mutations or their inverse activity in samples harboring established LOF mutations. **(f) Mutation heatmap without k-nearest neighbors correction.** When the analysis is performed by comparing PIK3CA^Mut^ samples to the centroid of *PIK3CA*^WT0^ samples rather than to their five nearest neighbors, stratification is dramatically degraded.

### Mutation Selection

For all subsequent analyses single point mutations and indels were downloaded from the TCGA firehose resource (gdac.broadinstitute.org, 2016-01-28 release http://firebrowse.org/), cBioPortal^17, 18^ and the GDC^29^ (see STAR Methods: *Mutations Selection*). Furthermore, a high-confidence GOF/LOF gold standard set was assembled by selecting literature-curated hypermorph and hypomorph mutations (**Figure 1b**). Assembly of this mutational gold standard was performed by literature datamining of the OncoKB^30^ and COSMIC^31^ databases, which also incorporate FASMIC^7^, using ProtFus^32^. All mutations (including both recurrent and rare ones) were included in the analysis. Samples harboring mutations with no prior experimental or established functional evidence were assigned to the unknown set.

### Proof-of-Concept Analysis

To test the proposed methodology, we first focused on *PIK3CA* and *TP53* as relevant case studies for GOF and LOF mutation assessment, because these genes present with a large number of both established and uncharacterized mutations, across multiple cancer cohorts (**Suppl. Figure 1b**).

*PIK3CA* and *TP53* Consensus Signatures were generated as described above using all established GOF and LOF events for these genes, in each tumor cohort presenting ≥ 5 such events. Notably, there were no reported LOF events for PIK3CA or GOF events for *TP53*. For each selected tumor cohort, all *PIK3CA* and *TP53* mutations of unknown significance were analyzed by PHNToM, while established mutations were re-classified by PHNToM by 5-fold cross validation. **Figure 2d** shows all samples harboring *PIK3CA* mutations in BRCA—including both functionally characterized events and events of unknown significance—sorted left to right from the one predicted as the most statistically significant GOF event to the one predicted as the most significant LOF event. Each row represents the activity of one of the 50 Consensus Signature TFs, as assessed by VIPER analysis of each sample vs. its P<u>^WT^</u> controls.

We performed a comprehensive analysis of *PIK3CA* across 25 TCGA cohorts, see **Suppl. Table 3.** We show heatmaps for the three tumor cohorts (UCEC, COAD, and BLCA) with the highest fraction of *PIK3CA*^Mut^ samples (49%, 29%, and 21%, respectively) (**Suppl. Figure 2a-c**). Similar to breast cancer, UCEC—which also represents a hormonally sensitive tumor—was predicted to harbor both GOF and LOF mutations, while the other three cohorts were predicted to harbor only GOF mutations. Similarly, we show heatmaps for four cohorts (BRCA, COAD, BLCA, and ESCA) with the highest fraction of *TP53*^Mut^ samples (52%, 51%, 42%, and 34%, respectively) (**Suppl. Figure 2d-g**). As shown across these heatmaps, the differential activity of Consensus Signature TFs was remarkably conserved across all established and predicted events and virtually perfectly inverted in LOF vs. GOF events, suggesting that these signatures may represent effective, highly conserved reporters of mutational activity.

Several observations emerge from these analyses: First, established *PIK3CA* GOF and *TP53* LOF mutations were virtually all ranked among those with the highest statistical significance by out-of-box, cross-validation analysis. Second, a significant number of events of unknown relevance appear to have an effect comparable to or greater than established GOF and LOF events, which was later confirmed by experimental assays. For *TP53*, in particular, the vast majority of previously uncharacterized mutations were predicted as LOF events. However, third, a small repertoire of *PIK3CA* (**Suppl. Figure 2a**) and *TP53* mutations (**Suppl. Figure 2d-f**) were predicted as LOF and GOF events, respectively, even though no such events are reported in databases.

To assess the improvements associated with use of the k-nearest neighbor approach for negative control selection, we generated the BRCA *PIK3CA* heatmap by comparing each *PIK3CA^Mut^* sample to the centroid of all *PIK3CA*^WT0^ rather than *PIK3CA*<u>^WT^</u> samples (**Figure 2e**). Compared to **Figure 2d**, the analysis shows dramatically decreased classification performance and strong co-segregation with tumor subtype.

### Structural Domain analysis

Proteins have complex 3D structures, with several structurally stable domains separated by floppy inter-domain regions. We thus asked whether spatial location, including as defined by structural domains, was associated with the functional relevance of mutational events. To accomplish this goal, we performed two analyses. First, we generated distinct sets of domain/interdomain-specific PIK3CA GOF mutations and compared their average behavior to the consensus signature generated from all PIK3CA GOF mutations. Second, we superimposed all established and statistically significantly predicted GOF mutations on the 3D structure of the protein, as recovered from the PDB database^33^. A 3D rendering of the spatial location of several established and predicted PIK3CA GOF mutations on the PDB 3D structure of the protein is shown in **Figure 3a**. This figure illustrates the spatial clustering of functionally relevant mutations in adjacent regions of the protein, a fact that cannot be captured by one-dimensional domain analyses, as also shown in^34^. *PIK3CA* mutations occur across all four major *PIK3CA* domains—namely, p85B, Ka, C2, and P14—as well as less frequently in inter domain regions^35^ (**Figure 3b, Suppl. Tables 4-5**). Most established GOF mutations in PIK3CA occur in the P14 (64 samples) and C2 (42 samples) domains. Consistent with this, established GOF mutations in these two domains provided the best and second-best match to the global *PIK3CA* Consensus Signature **(Figure 3c)**. Indeed, mutations in the P14 domain presented even more statistically significant differential activity of Consensus Signature TFs. Consistent with these findings, VIPER-based assessment of PIK3CA activity in BRCA was significantly higher for samples harboring mutations in these two domains compared to both PIK3CA^WT^ samples and normal breast epithelial samples in GTEx^36^ **(Figure 1b)**. This also suggests that domains other than P14 and C2 are more likely to be enriched in neomorphic events. Indeed, most established *PIK3CA* neomorphic mutations^6^ (*e.g., PIK3CA^E545^, PIK3CA^E542^, PIK3CA^H1047^, PIK3CA^Q546K^*) occur outside of the P14 and C2 domains.

**Figure 3:**
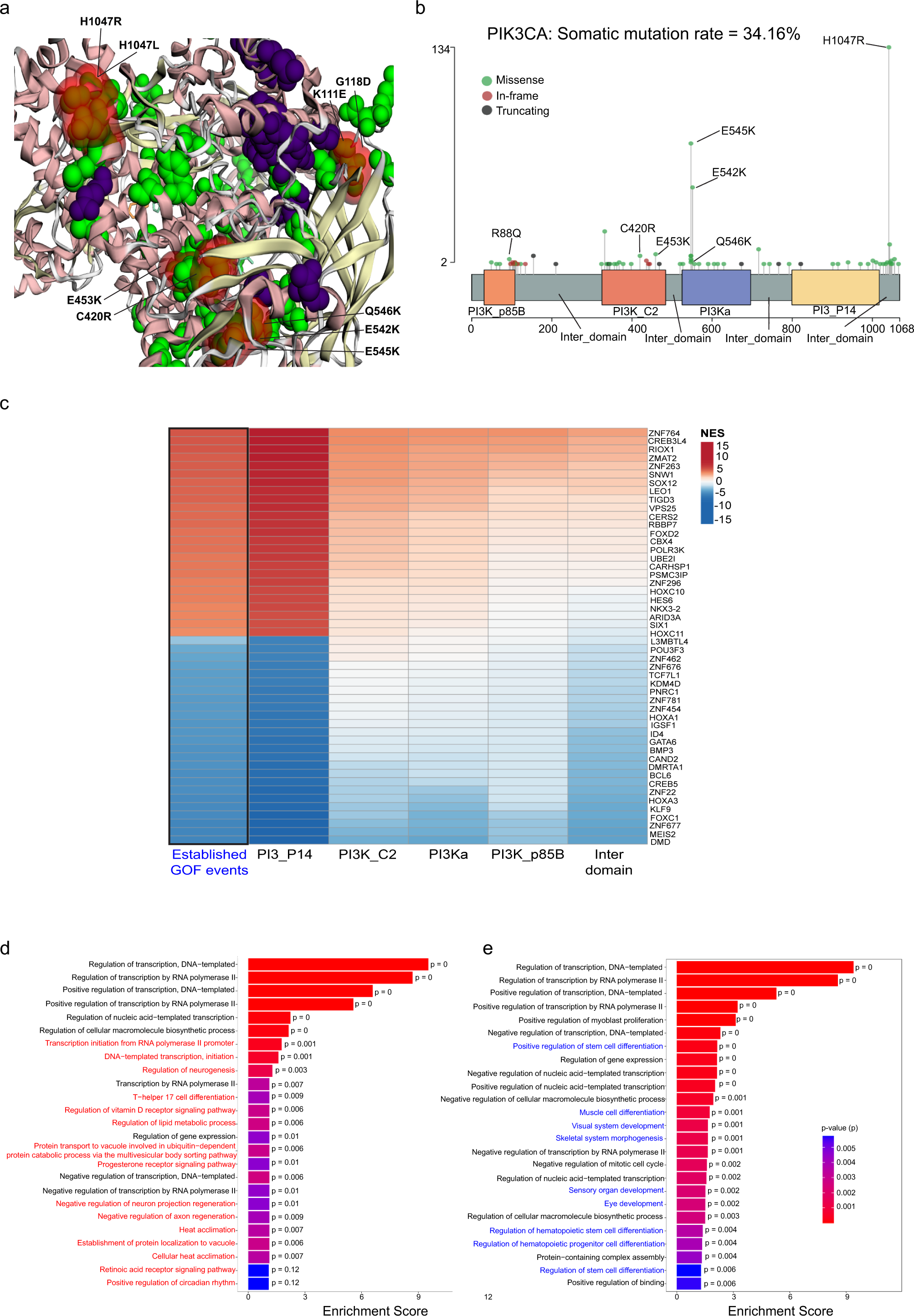
Domain-wise analysis of PIK3CA mutations in BRCA. **(a)** This panel shows the spatial location of both established and predicted GOF PIK3CA mutations on the PDB 3D structure of the protein. Clustering of functionally relevant mutations in spatially-adjacent region of the protein is evident. **(b)** This shows the occurrence of missense, in-frame shifts, and truncating mutations in distinct PIK3CA protein domains—including PI3K_p85B, PI3K_C2, PI3Ka, PI3_P14—as well as occurring in inter-domain regions. **(c)** The first column shows the activity of TF/co-TF proteins in the Consensus GOF/LOF Signature, averaged across all samples harboring GOF events. The following columns show differential TF/co-TF activity in GOF events, averaged on a domain-by-domain basis. **(d-e)** Enrichment analysis of Gene Ontology and Reactome functional categories in the 50 most activated and 50 most inactivated TF/co-TF proteins, respectively. Categories that are differentially enriched in the two analyses are shown in red and blue, respectively.

### Mutational Effector Analysis

The striking consistency of differentially activated and inactivated TF/co-TF proteins in PIK3CA^Mut^ vs. PIK3CA<u>^WT^</u> samples further supports their potential as conserved downstream effectors of *PIK3CA* activity (**Figure 3c**, first column), including on a domain-by-domain level (other columns). In addition, the most aberrantly activated and inactivated TFs/co-TFs presented key differences in terms of Gene Ontology and Reactome functional category enrichment **(Figure 3c** and **d, respectively).**

Some established PIK3CA GOF mutations were predicted to have significantly stronger effects than others. For instance, compared to PIK3CA^H1047L^ and PIK3CA^D603H^, PIK3CA^N345K^ and PIK3CA^E726K^ produced significantly higher VIPER-inferred PIK3CA activity and PHNToM scores **(Figure 4a)**. This may help stratify patients that are most likely to respond to a PIK3CA pathway inhibitor as well as to tailor the use of each inhibitor based on the ability to reverse the specific activity pattern induced by the mutation (OncoTreat algorithm). However, more importantly, a large number of novel, PHNToM-predicted mutations had GOF scores as high or higher than the top established mutations (**Figure 4a**, red proteins) (**Suppl. Table 6**). In addition, PHNToM identified mutations producing an almost perfect Consensus Signature activity inversion, thus most likely to represent LOF events, even though no PIK3CA LOF mutations were previously reported (**Suppl. Table 6**). Finally, the analysis identified a significant number of likely neutral mutations (**Suppl. Table 6**). Prediction of NEU and LOF event is equally important because patients harboring these mutations would almost unequivocally fail to benefit from PI3K inhibitors.

**Figure 4:**
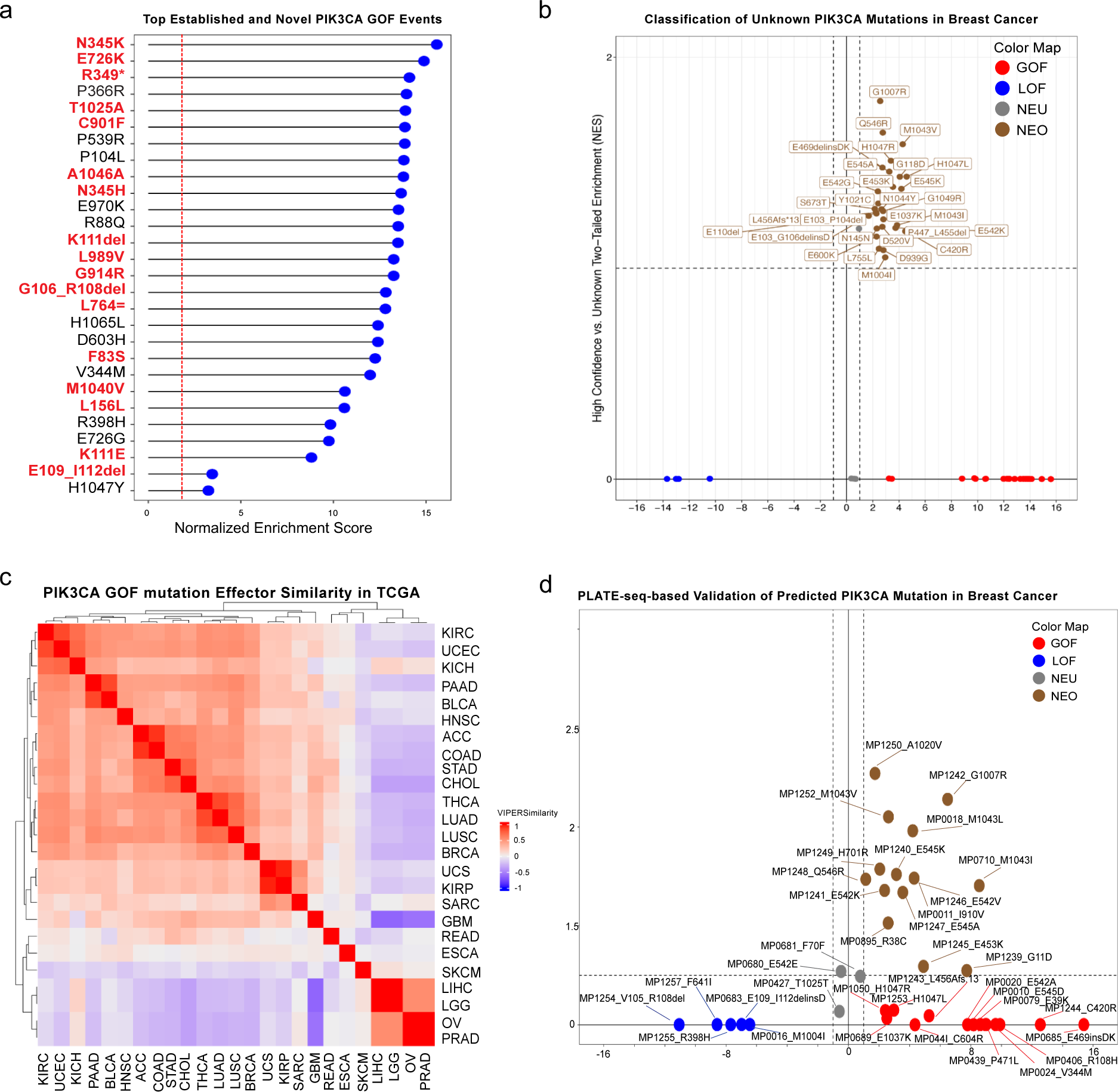
PIK3CA Mutational Analysis. **(a)** Established (black) and predicted (red) GOF mutations are shown, ranked by the average of their DGN and IGN. Novel, predicted GOF events have strength comparable or greater than established ones. **(c)** Scatter plot showing all PIK3CA mutations of unknown significance identified as non-neutral by PHNToM in the BRCA cohort, including GOF (red), LOF (blue), NEU (gray), and NEO (brown). **(b)** Conservation of PIK3CA effector TF/co-TF proteins across 25 TCGA cohorts, using the OncoMatch similarity metric, reveals two main clusters sharing similar consensus signatures**. (d)** Validation of predicted GOF, LOF, NEO, and NEU events based on DGN and IGN analysis of MCF10 cells following mutation knock-in vs. non-targeting controls, as generated by PLATE-seq gene expression profiles.

To optimally visualize these findings, we use a 2d scatter plot, with × = - Log *p(GOF; LOF)* and *y = - Log p(s_NEO_*), see **Figure 4b** for all statistically significant PIK3CA mutations in BRCA. Bonferroni corrected adjusted p-values associated with all these assessments are provided in **Suppl. Table 6**. High-confidence, PHNToM-predicted neomorphic events and associated TCGA sample accession number are included in **Suppl. Table 7-9**.

### Pan cancer analysis

When extended to all 25 TCGA cohorts, the PIK3CA analysis identified novel GOF, LOF, NEO, and neutral mutations in 13 cohorts (**Table 1** and **Suppl. Table 3**).

**Table 1:**
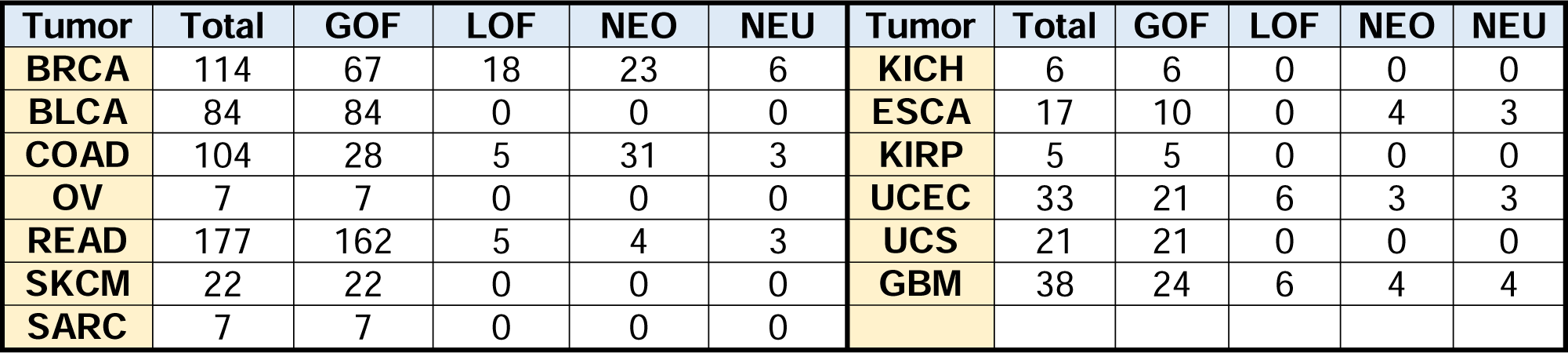
PIK3CA mutation characterization across relevant TCGA cohorts.

| Tumor | Total | GOF | LOF | NEO | NEU | Tumor | Total | GOF | LOF | NEO | NEU |
| --- | --- | --- | --- | --- | --- | --- | --- | --- | --- | --- | --- |
| BRCA | 114 | 67 | 18 | 23 | 6 | KICH | 6 | 6 | 0 | 0 | 0 |
| BLCA | 84 | 84 | 0 | 0 | 0 | ESCA | 17 | 10 | 0 | 4 | 3 |
| COAD | 104 | 28 | 5 | 31 | 3 | KIRP | 5 | 5 | 0 | 0 | 0 |
| OV | 7 | 7 | 0 | 0 | 0 | UCEC | 33 | 21 | 6 | 3 | 3 |
| READ | 177 | 162 | 5 | 4 | 3 | UCS | 21 | 21 | 0 | 0 | 0 |
| SKCM | 22 | 22 | 0 | 0 | 0 | GBM | 38 | 24 | 6 | 4 | 4 |
| SARC | 7 | 7 | 0 | 0 | 0 |  |  |  |  |  |  |

PIK3CA mutations were uncommon in other cohorts, thus precluding effective analysis. *PIK3CA* mutation classification plots for selected cohorts with the greatest fraction of mutated samples, including UCEC, COAD, BLCA, and ESCA, are shown in **Suppl. Figure 3a-e.**

A key question is whether PIK3CA effectors are conserved across distinct tumor types. Based on GOF/LOF Consensus Signature similarity by Spearman Correlation, two clusters emerged with virtually orthogonal effector proteins (**Figure 4c**). The first cluster was comprised almost exclusively of epithelial tumors, while the second one comprised PRAD, OV, LGG, SKCM, and LIHC. In contrast, stomach adenocarcinoma (ESCA) had highly unique effector signatures that did not match any other tumor. The same was observed for *TP53*^Mut^, with a virtually identical set of cohorts where the consensus signature was conserved or discordant, albeit with KIRP, SARC, GBM, and READ, forming a third cluster and ESCA still representing an outlier (**Suppl. Figure 2h**). Taken together, these data suggest that PIK3CA and p53 may regulate distinct TF/co-TF effectors in different tumor types likely as a consequence of lineage specific gene expression patterns.

### Retrospective Validation

Analysis of published data (FASMIC) on neomorphic activity, produced a statistically significant contingency table comprising 18 True Positive (TP), 15 True Negative (TN), 5 False Positive (FP), and no False Negative (FN) predictions (p = 1.1×10^−^06, by χ^2^ analysis) **(Suppl. Table 10-11)**. Note that statistical significance is likely underestimated due to lack of systematic validation.

Despite the fact that neomorphic events have not been systematically assessed, comprehensive literature survey shows that some of the neomorphic events predicted by PHNToM were previously reported. For instance, *PIK3CA^E545^, PIK3CA^E542^, PIK3CA^H1047^, PIK3CA^Q546K^* and *PIK3CA^G1049R^* were characterized in^6^. Similarly, mass spectrometry *analysis showed that PIK3CA^E545K^* induces an unexpected interaction with insulin receptor substrate 1 (*IRS1*), independent of *p85*^37^.

Supporting the ability to identify tissue-specific hypermorphs, PHNToM predicted the missense mutation *PIK3CA^V71I^* as GOF in a single Head and Neck Squamous Cell Carcinoma (HNSC) sample (*TCGA.T2.A6WZ)* (NES = 15.97) but not in BRCA. Consistent with this prediction, FASMIC^3^, reported this mutation as GOF in H&N cells but not in the breast epithelial cell line MCF10^38^.

### Prospective Validation of PHNToM predictions in MCF10 cells

To prospectively validate PHNToM predictions, we selected 37 breast cancer-specific *PIK3CA* mutations, comprising 9 GOF, 5 LOF, and 20 NEO PHNToM-predicted events, whose functional relevance had not been previously assessed in FASMIC. For this purpose, differential gene expression signatures were generated from MCF10 cell, following lentivirus-mediated knock-in of mutant PIK3CA cDNA vs. mock cDNA as negative control. RNA-seq profiles were generated in 96-well plates by Pooled Library Amplification for Transcriptome Expression (PLATE-Seq)^16^, a highly controlled microfluidic assays. Briefly, PLATE-Seq introduces well-specific barcodes during reverse transcription, thus supports pooled library construction and low-depth sequencing with an average of 2M reads/sample. Functional activity was then assessed based on the ability of each knock-in to recapitulate the human breast cancer-specific *PIK3CA* Consensus Signature by DGS and IGS enrichment analysis.

The analysis confirmed predictions for 15/15 GOF (100%), 5/5 LOF (100%), 15/20 NEO (75%) and 3/3 NEU variants (100%) (p < 1e-4, by two-tail χ^2^, see methods) (**Figure 4d**), with individual predictions assessed at p ≤ 0.05 (Benjamini-Hochberg corrected). Expected outcome probabilities were computed based on the fraction of established GOF, LOF, NEO, and NEU PIK3CA mutations in FASMIC. Bonferroni corrected adjusted p-values associated with each event are provided in **Suppl. Table 12**. PAEA plots for some validated GOF, LOF and NEO events are shown in **Figure 5a-c**. Even though no known *PIK3CA* LOF mutations have been reported in the literature, PHNToM-predicted LOF mutations were confirmed by these assays.

**Figure 5:**
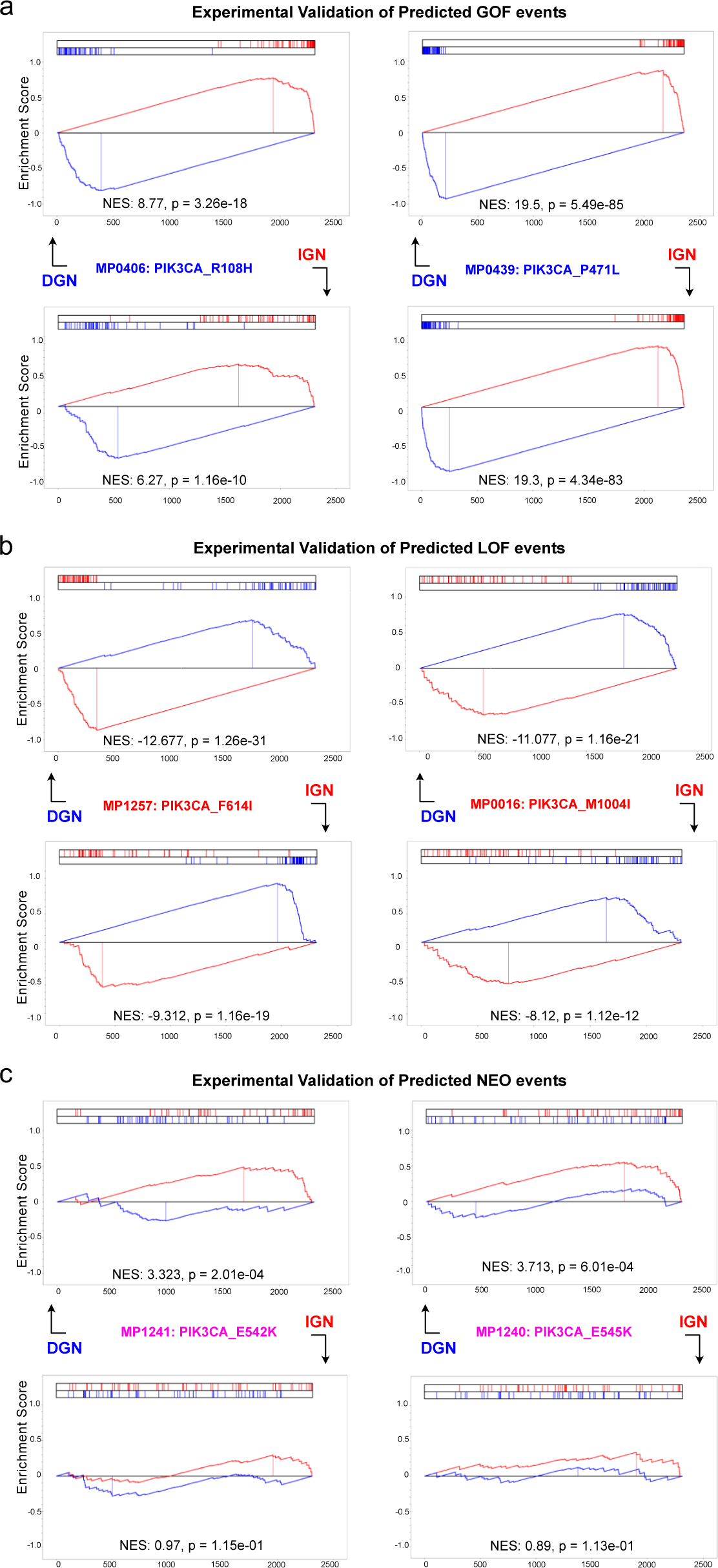
Illustrative Example of DGN and IGN analysis of PLATE-seq profiles generated by knock-in of PHNToM-predicted GOF, LOF, and NEO mutations in MCF10 cells. **(a)** Validation of *PIK3CA*^R108H^ and *PIK3CA*^P471L^ as GOF events. **(b)** Validation of *PIK3CA*^F614I^ and *PIK3CA*^M1004I^ as LOF events. **(c)** Validation of *PIK3CA*^E542K^ and *PIK3CA*^E545K^ as NEO events.

### Systematic Analysis of TCGA Cohorts

For each of the 25 TCGA cohorts with ≥ 100 samples (**Suppl. Figure 1a, Suppl. Table 1**), thus supporting ARACNe-based network assembly, we used PHNToM to analyze every gene for which established GOF and/or LOF mutations were reported in the GOF/LOF gold standard set. To avoid deriving idiosyncratic GOF/LOF Consensus Signatures, due to potential co-mutational events, we restricted the analysis only to genes for which 5 or more samples harboring established GOF/LOF were available. Consensus Signatures for each gene/tumor-cohort pair satisfying these constraints were then generated, as described in the previous sections, and used to characterize all other events of unknown functional significance. Overall, 3,736 genes could be analyzed in at least one TCGA cohort, see **Suppl. Table 1** for a detailed report of protein/cohort pairs and for the TCGA IDs of associated, mutation-harboring samples.

Due to space limitations, we restrict the report of detailed functional predictions to the 20 most recurrently mutated genes in cancer—namely, TP53, PIK3CA, KMT2D, FAT4, ARID1A, PTEN, KMT2C, APC, KRAS, BRAF, FAT1, ATRX, NF1, IDH1, ATM, ERBB2, FGFR2, ALK, KDR, RET, PIK3R1, and GNAS **(Suppl. Table 3)**. For each mutation, we show the event’s location on the protein structure, as well as any literature evidence, frequency, functional prediction and dysregulated protein activity, compared to its P<u>^WT^</u> controls. A more comprehensive analysis of all mutation-bearing proteins will be provided on a dedicated website upon publication.

### Mutational Mimicry Assessment

Mutations in two different genes can phenocopy each other’s functional effects, as assessed based on the differential activity of the downstream TF effectors identified by PHNToM analysis. This is consistent with the concept of pseudo-mutants^39^ or mutational mimicry^40^. While this is not unreasonable for genes encoding proteins in the same pathway, such as different MAP kinases, detection of mimicry in more distal genes may underlie unknown mechanisms of regulatory/signaling interactions, canalization, and cross-talk.

For instance, consistent with the fact that p53 regulates *PIK3CA* on its promoter, PHNToM shows that samples with *PIK3CA* GOF mutations are highly enriched in the LOF consensus signature for *TP53* (**Figure 6a**), suggesting that *TP53* LOF mutations mimic the effect of *PIK3CA* LOF mutations. This provides a potential mechanistic rationale to interpret some otherwise puzzling results. For instance, consistent with the literature, PHNToM classifies *PIK3CA^E545K^* as a neomorphic event in BRCA. However, it also classifies this mutation as an hypomorphic event in the BRCA Luminal A sample*TCGA.EW.A1J5*. While this could be simply a false positive prediction, a more likely explanation is that, consistent with **Figure 6a**, this sample also harbors a *TP53^X331_splice^* LOF mutation, which, as discussed phenocopies the effect of a *PIK3CA* LOF mutation by decreasing the expression of the gene.

**Figure 6:**
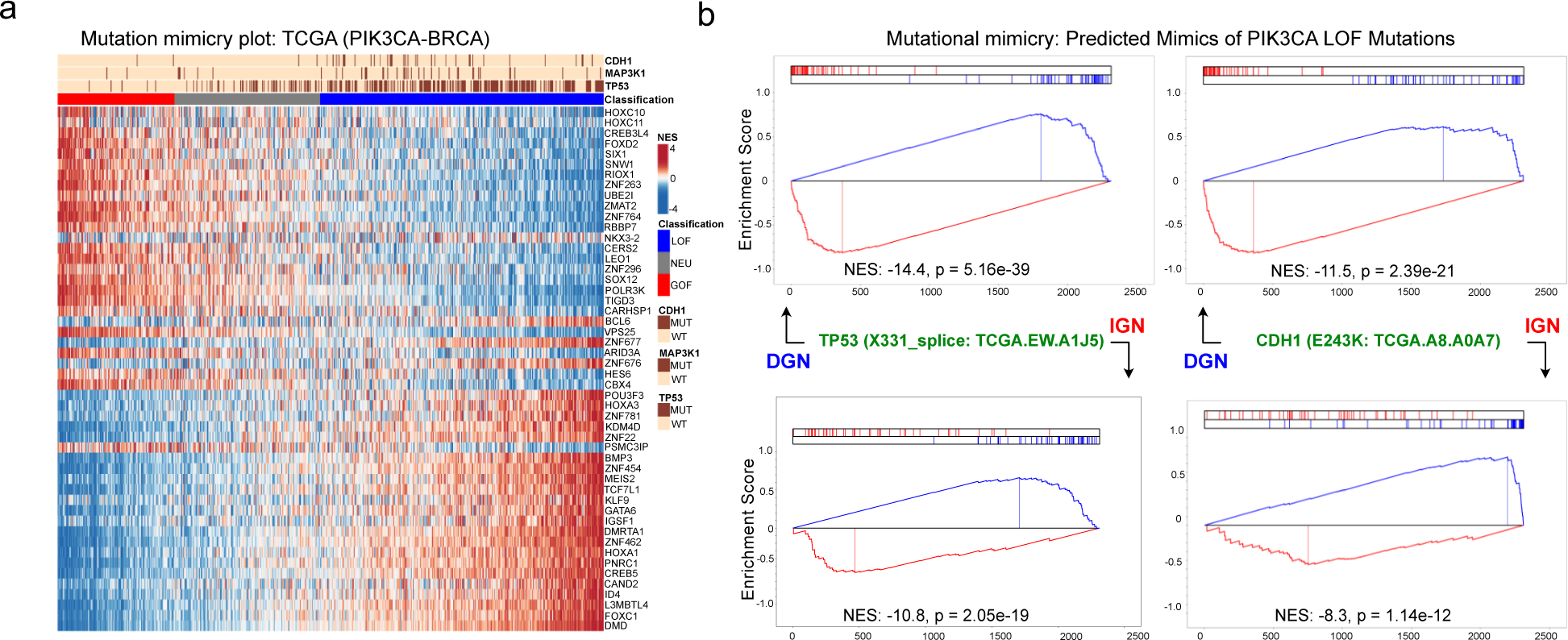
Mutational mimicry analysis. **(a)** A mimicry heatmap for PIK3CA LOF mutations shows that BRCA samples harboring TP53 LOF mutations are dramatically enriched in samples presenting a negative *PIK3CA* GOF/LOF Consensus Signature, suggesting that *TP53* LOF events induce PIK3CA LOF. *CDH1* and *MAP3K1* LOF mutations induce similar effects, albeit resulting in more modest PIK3CA inactivation. **(b)** DGN and IGN enrichment analysis for illustrative mutations in *TP53* and *CDH1*.

By leveraging the activity of Consensus Signature TFs as a high-dimensional reporter assay, we generalized mimicry from a few anecdotic examples to a comprehensive map of interactions. This map may help interpret the effect of GOF/LOF mutations that fail to induce their expected effects in specific tumors—including providing a rationale for targeted therapy failures—as well as the complementarity between different genetic alterations. In addition, it may help extend use of available inhibitors that target downstream events to mutational mimics of their target genes.

Systematic analysis of all possible recurrently mutated gene pairs, across all 25 TCGA cohorts, is summarized in Suppl. Table 3. Specifically, for each gene, we report all other mutations that are associated with either positive or negative enrichment in its GOF/LOF Consensus Signature. We also represented these findings as a functional interaction heatmaps, see **Suppl Figure 4** for a mimicry map of ARID1A LOF in UCEC, including the effect of KRAS, EGFR, and KDR mutations, with associated DGN and IGN enrichment plots.

## Discussion

In this manuscript, we leverage the activity of all transcriptional regulator proteins as a highly-multiplexed reporter assay to computationally elucidate the function of specific mutations as hyper (GOF), hypo (LOF), neomorphic (NEO) or neutral events, via the PHNToM algorithm. Experimental validation of uncharacterized mutations, predicted to be functionally relevant by the analysis, and retrospective benchmarks for previously characterized events, suggest that this approach can effectively complement experimental methods, without requiring the expensive, labor-intensive protocols necessary for tumor context-specific functional characterization.

While most commonly occurring mutations have been reported as either GOF or LOF, our analysis complements previous reports, see^34^ for instance, in suggesting the pervasive nature of neo-morphic mutations in oncogenes and tumor suppressors, ranging from *PIK3CA* and *EGFR* to *PTEN* and *TP53*. As also shown^41^, some neomorphic mutations could be explained as inducing substantial rewiring of signaling networks, that are insensitive to feedback loops, resulting in hyperactivation that is sufficient to activate mechanisms not normally controlled by their WT counterpart. This may be further reflected by the fact that several of the neomorphic mutations identified by the analysis also presented slight yet statistically significant hyper or hypo-morphic activity in addition to their apparent neomorphic activities. Regardless of the underlying mechanisms, this underscores the importance of distinguishing between these three classes of functionally distinct mutations, including a need to identify their suitability for treatment with targeted inhibitors. Just as important, accurate characterization of passenger mutations that do not affect the activity of the protein may also significantly improve use of targeted agents by preventing their use in patients who would not benefit.

Notably, our analysis showed that most neo-morphic mutations in oncogenes comprise either substitution or truncation mutations. For example, in *PIK3CA*, the majority of neomorphic mutations cluster in three hotspots: *PIK3CA^E542^* and *PIK3CA^E545^* in the helical domain encoded by exon 9 and *PIK3CA^H1047^* on exon 20 which encodes for the kinase domain. *PIK3CA^E542^* and *PIK3CA^E545^* are commonly substituted with a lysine while *PIK3CA^H1047^* is often changed to an arginine. These mutants were originally classified as hyper-morphs due to their altered interaction with p85 or to conformational changes of the activation loop, leading to constitutive low-level activity of the *PIK3CA^E542^* and *PIK3CA^E545^* variants^38^ and increased sensitivity of the *PIK3CA^H1047^* mutant to ligand activation, resulting in increased tumorigenicity^38^. Since these recurrent *PIK3CA* mutations are present in a large number of cancer patients, understanding the neomorphic functions of the helical domain mutants is critical to effectively capitalize on the promise of targeted therapy. Indeed, neomorphic activity of these variants may substantially contribute to the limited activity of PI3K and pathway inhibitors in *PIK3CA* mutant tumors. Such a mechanism was previously suggested based on the study of a hydrocarbon-stapled mutant peptide that disrupted the neo-morphic interaction between IRS-1 and the *PIK3CA^E545K^* mutant^42^. Strikingly, this peptide inhibited AKT phosphorylation and tumor growth in xenografts with the *PIK3CA^E545K^*neomorph but not in those with other mutations. This suggests potential therapeutic approaches leveraging targeted agents capable of interfering with a protein’s neomorphic activity without affecting its WT function, potentially resulting in a wider therapeutic index. In addition, small molecule inhibitors targeting signaling pathways downstream of *PIK3CA,* or activated by *PIK3CA* neomorphs, could also represent potential therapeutic opportunities.

Our study also illustrates the importance of assessing the functional effects of specific mutations in a tumor context-specific manner, including within distinct sub-lineages of the same tumor type. Indeed, analysis of the same mutational event, across multiple cohorts, can identify it as hypomorphic, hypermorphic or neomorphic in some but not others, likely due to context-specific availability of cognate binding partners and epigenetically-mediated activation or silencing of critical downstream targets. For instance, *PIK3CA^R108H^, PIK3CA^G118D^* and, *PIK3CA^E110del^* were predicted to be neomorphic in BRCA yet hypermorphic in UCEC.

Finally, while a few anecdotic cases had been previously reported, our study identified an extensive repertoire of molecular mimics that phenocopy the effect of mutations in unrelated proteins, based on the transcriptional effectors identified by the PHNToM analysis. These findings can be critical to identify signaling pathway crosstalk and novel molecular interactions. For instance, in UCEC, in *EGFR^A743T^* (TCGA.A5.A0G2), *KRAS^Q61H^* (TCGA.A5.A0G2), *KDR^S1347L^* (TCGA.A5.A0G2) mutations almost perfectly recapitulate the effect of *ARID1A* LOF mutations.

Overall, our analysis was able to characterize more than half million mutational events that are still uncharacterized in public databases, thus massively improving our knowledge on the functional nature of oncogene mutations. Interestingly, while the validation rate for neomorphic mutations was a bit lower (75%) compared to GOF and LOF mutations—likely because their effect may be difficult to detect/recapitulate in cell lines that may have slightly altered cognate binding partner repertoires compared to tumors—the validation rate for GOF and LOF was virtually perfect (100%), albeit based on a relatively small set of tested mutations. This suggests that PHNToM provides a valuable and highly complementary approach to the accurate assessment of mutational events, with potential applications to targeted therapy selection.

## Supplemental Figures

***Suppl. Figure 1: PIK3CA and TP53 mutations in TCGA cohorts.* (a, b)** Histograms of the number of samples in each TCGA cohorts harboring *PIK3CA* and *TP53* mutations, respectively.

***Suppl Figure 2****: **PIK3CA and TP53 Mutation heatmaps for selected TCGA cohorts*****. (a-c)** Protein activity heatmaps for PIK3CA Consensus Signature TFs for UCEC, COAD and BLCA cohorts. **(d-g)** Protein activity heatmaps for TP53 Consensus Signature TFs for UCEC, COAD and BLCA cohorts. **(h) Correlation analysis.** Clustered heatmap showing the correlation of TP53 Consensus Signature TFs across all 25 TCGA cohorts, as assessed by OncoMatch.

*Suppl. Figure 3: Mutation classification plots for PHNToM-predicted GOF, LOF, NEU, and NEO events in different TCGA cohorts.* (a-d) Similar to Figure 4b, these scatter plots stratify the effect of PIK3CA mutations based on the activity of Consensus Signature TFs. The x-axis represents the statistical significance (-Log p) of the DGN + IGN metric, representing the GOF/LOF effect, while the y-axis represents the statistical significance of the |DGN – IGN| metric, representing the NEO effect. Different panels correspond to analyses in the UCEC, COAD, BLCA, and ESCA cohorts. **(e)** Conservation of BRCA Consensus Signature TFs in the Consensus Signature of the UCEC, COAD, BLCA, and ESCA cohorts. Consistent with Figure 4c, conservation is highly statistically significant for the first three cohorts but not for ESCA.

***Suppl Figure 4****: **Mutational mimicry heatmap for ARID1A in UCEC*****. (a)** Mimicry analysis shows that mutations in KDR, EGFR, and KRAS are significantly enriched in samples presenting an ARID1A GOF Consensus Signature. (b – d) DGN and IGN analysis representing the enrichment of the most differentially active TF/co-TF proteins in samples harboring *EGFR*^A743T^, *KRAS*^Q61H^, and *KDR*^S1347L^ mutations in the ARID1A Consensus Signature (DGN) and vice-versa (IGN).

## ACKNOWLEDGEMENTS

G.B.M. is supported by NCI grants U24 CA 264128, U01 CA253472, U01 CA 217842 and U01 CA 281902 and a kind gift from the Dr. Miriam & Sheldon G. Adelson Medical Research Foundation. A.C. was supported by the NCI Cancer Target Discovery and Development Program (U01 CA217858 and U01 CA272610) and NIH Shared Instrumentation Grants (S10 OD012351 and S10 OD021764).

## AUTHOR CONTRIBUTIONS

So.T., G.B.M., and A.C. conceived this work. So.T. designed, performed, and oversaw the computational analyses. Sa.T. performed experiments. So.T., G.B.M. and A.C. wrote the manuscript. So.T., Sa.T., G.B.M. and A.C. reviewed the manuscript. All authors approved the final manuscript.

## CODE AVAILABILITY

Full code is available at: https://github.com/somnathtagore/PHaNToM

## STAR Methods

### TCGA sample selection and normalization

RNA-Seq profiles comprising raw, paired-end reads from 25 TCGA cohorts were downloaded from Genomic Data Commons Data Portal (GDC, https://portal.gdc.cancer.gov/, v15.0, Release 2019/02/20)^29^. Equivalent profiles for GTEx samples were downloaded from The database of Genotypes and Phenotypes (dbGaP, https://dbgap.ncbi.nlm.nih.gov/aa/wga.cgi?page=login)^43^ and Genotype-Tissue Expression (GTEx, https://gtexportal.org/home/, GTEx Analysis V7, dbGaP, Accession: phs000424.v7.p2)^36^. STAR Aligner (v 2.6.1e 2019/07/25)^44^, was used to map individual reads to the UCSC^45^ human reference genome hg38 (patch release 13, GRCh38.p13, 2019/02/28) and to the reference transcriptome GENCODE (v29, GRCh38.p12, 2017/12/21), followed by FeatureCounts (Release 2.0.0, 2019/09/04)^46^ analysis to generate read counts.

To correct for batch effects, independent sample-gene matrix were created for each TCGA tumor cohort and tissue-match normal GTEx tissue pair. ComBat (version 3.27.0)^47^ was used to batch-correct for technical bias between TCGA and GTEx. RNA-Seq raw gene counts were then transformed to Reads Per Kilobase of transcript, per Million mapped reads (RPKM) using the average transcript length for each gene, followed by log2 transformation.

### ARACNe network generation

Gene expression signatures were normalized by applying two non-parametric transformations for ARACNe analysis. This is done because mutual information estimation is not affected by monotonic transformation of input data. Specifically, rank transformation between 0 and 1 was performed, first on a column basis (tumor sample) and then again on a row basis (gene). The resulting data was analyzed by ARACNe-AP^22, 48^ to generate tissue specific regulons for each TF and co-TF protein.

### VIPER analysis

VIPER was used to transform gene expression profiles into protein activity profiles for all TF and co-TF proteins, on a sample-by-sample basis. Specifically, for each sample (either tumor or normal) the differential expression signature obtained by Student T-test analysis of genes in each sample vs. the average of all samples in the GTEx tissue matched cohort was analyzed by VIPER algorithm (v1.24.0)^14^, using the tissue-matched ARACNE-AP regulons.

### Mutation selection

Mutation Annotation Files (MAF) for TCGA were downloaded from the firehose repository (gdac.broadinstitute.org, 2016-01-28 release) (http://firebrowse.org/). Additional mutational data were downloaded from cBioPortal (Release 2019/03/15; 2021/01/28)^17, 18^ and GDC (v15.0, Release 2019/02/20)^29^. Somatic copy-number alterations were assessed based on published GISTIC2.0 (v7) analyses^49^.

### P^WT0^ Null Model Generation

To identify a set of samples (P^WT0^) representing the unbiased activity of the wild-type protein (P^WT^), we generated a probability density from GTEx samples representing the VIPER-assessed activity of P in the most closely related normal tissue counterpart and then selected tumor samples with VIPER-assessed protein activity not statistically significantly different from that of the normal cohort (p > 0.5).

### Multi-modal Gaussian mixture model

For each protein under consideration, the pipeline performs a multi-modal analysis of WT, mutated, and GTEx samples (either tissue matched if available or averaged over the entire GTEx repository). Then, normal controls are selected as WT samples that are indistinguishable from the GTEx samples (i.e., p > 0.05, based on Student’s T-test).

### Multi-modal Gaussian mixture analysis

These analyses are based on probabilistic models used for representing normally distributed samples within a bigger sample space, parameterized by two types of values, the mixture component weights and the component means as-well-as variances/co-variances^50^. For a Gaussian mixture model with *C* components, the *c^th^* component has a mean of μ*_c_* and variance of σ*_c_* for univariate and mean 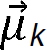 and covariance matrix of Σ*_k_* for multivariate. The mixture component weights are defined as □*_k_* for component *C_k_*, with the constraint that 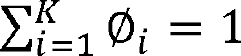 so that the total probability distribution normalizes to 1. If the component weights aren’t learned, they can be viewed as an a-priori distribution over components such that *p*(*x* generated by component *C_k_*)=□*_k_*. If they are instead learned, they are the a-posteriori estimates of the component probabilities given the data.

### K-nearest neighbor analysis

To remove confounding effects arising from the differential representation of specific mutations in transcriptionally distinct cancer patient subsets, we used a k-nearest neighbors^51^ approach with k = 5. Specifically, for each sample *S_i_* harboring a mutation of interest, negative controls were selected as its k-nearest neighbors among the subset of P^WT0^ samples. The number of samples (k = 5) was determined empirically to (a) provide reasonable statistics on differential expression and (b) increase the likelihood that negative controls were selected from the same genetic background subpopulation.

### Consensus GOF/LOF Signature

To build the consensus signature based on established hypermorph (GOF) or hypomorph (LOF) mutations, the algorithm was trained using the sample subsets defined in cBioPortal (Release 2019/03/15; 2021/01/28)^17, 18^, OncoKB (v1.24, 2019/12/12; v3.0, 2021/01/14)^30^, FASMIC (v1.8, 2021/11/01)^3^ or the literature. Specifically, vectors representing the differential activity of each TF in each sample harboring a GOF mutation, compared to its P<u>^WT0^</u> negative controls, were generated by VIPER analysis. For LOF samples, the same procedure was used with an inverted sign for protein activity, such that the most differentially active protein would become the most differentially inactive one. Finally, all GOF and LOF vectors were Stouffer integrated^52^ to generate a single vector of TF/co-TF proteins rank-sorted based on the statistical significance of their integrated differential activity, thus representing the consensus GOF/LOF signature.

### Null model to estimate the statistical significance of the PIK3CA validation assays

Statistical significance was computed by Χ^2^ analysis for GOF, LOF, NEO, and NEU events using a null model where the expected probability of validation was computed based on the ratio between the number of validated GOF, LOF, NEO, and NEU events and the total number of validated PIK3CA mutations in the cohort.

