## Supplementary figures and images for "Systematic Pan-cancer Functional Inference and Validation of Hyper, Hypo and Neomorphic Mutations"

### Suppl. Figure

Suppl. Fig. 1

b

*PIK3CA* mutations in TCGA cohorts

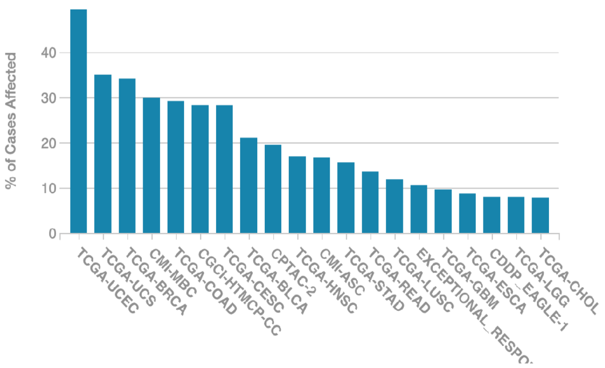


*TP53* mutations in TCGA cohorts

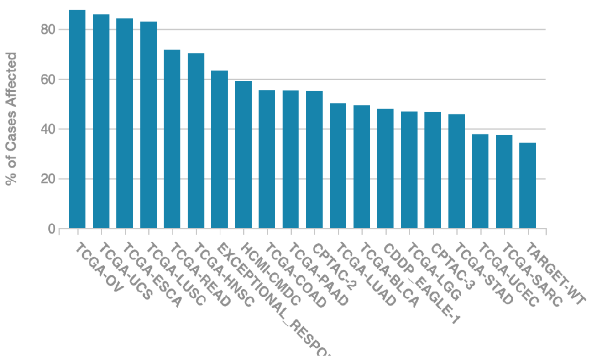

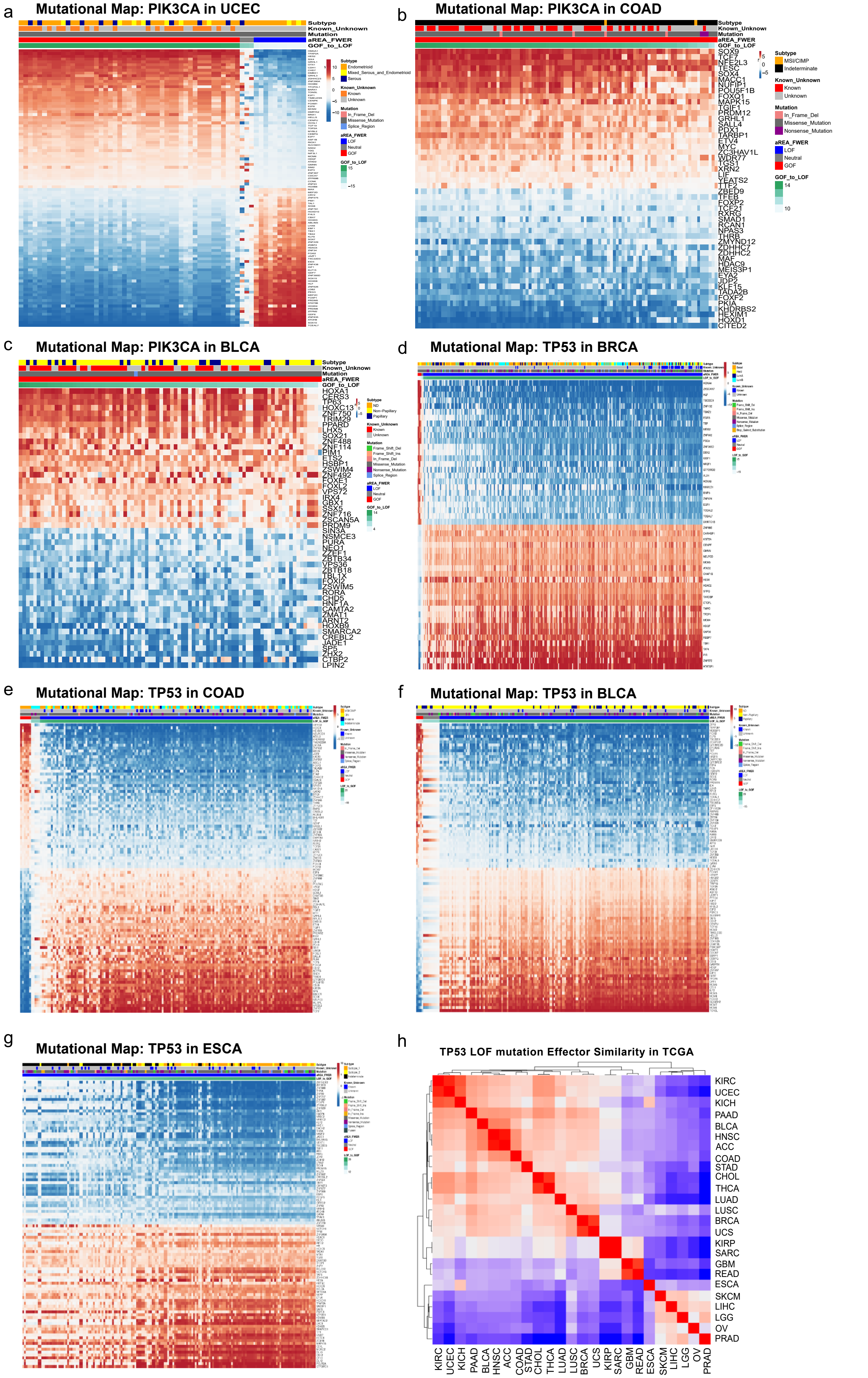

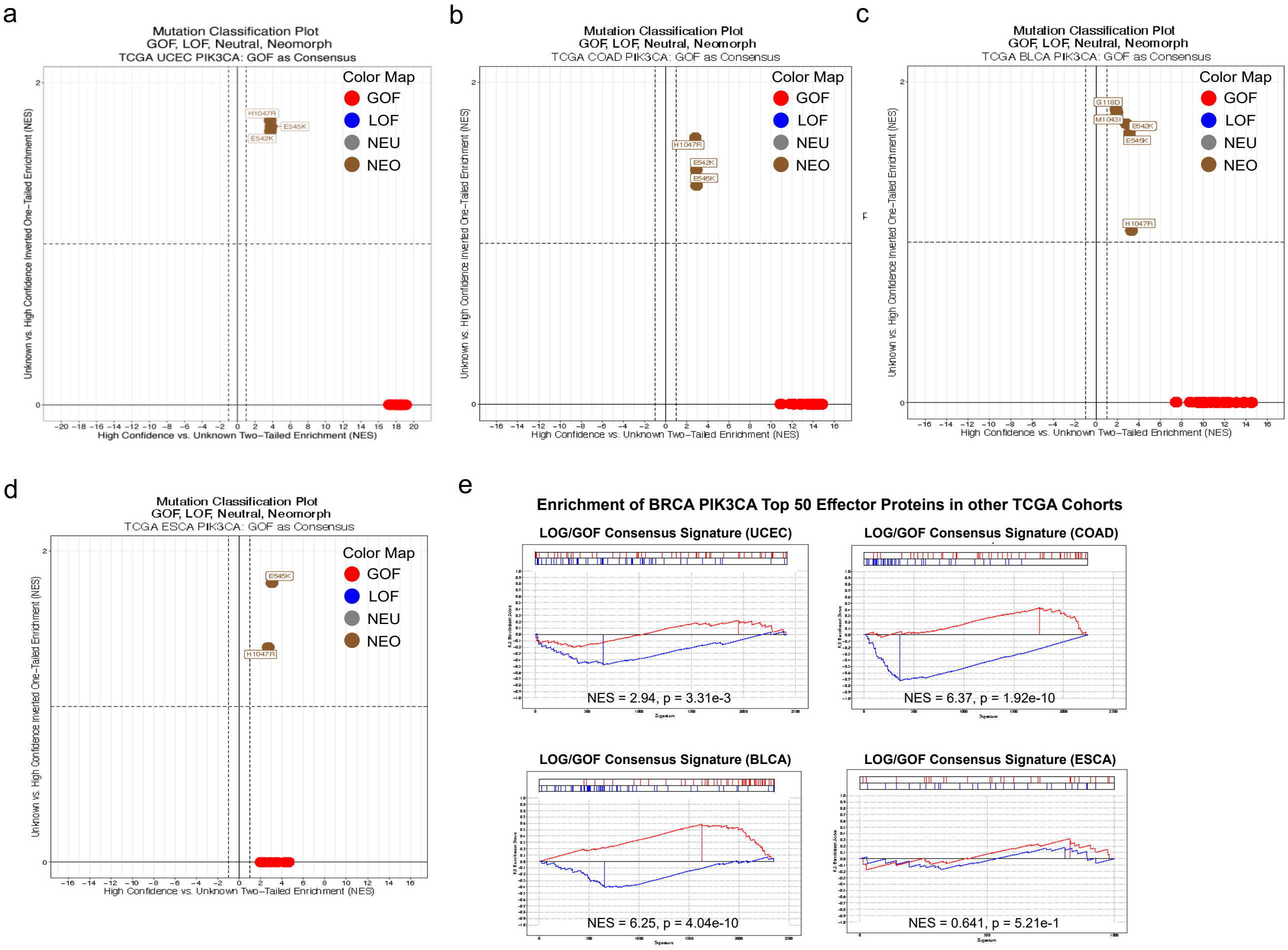

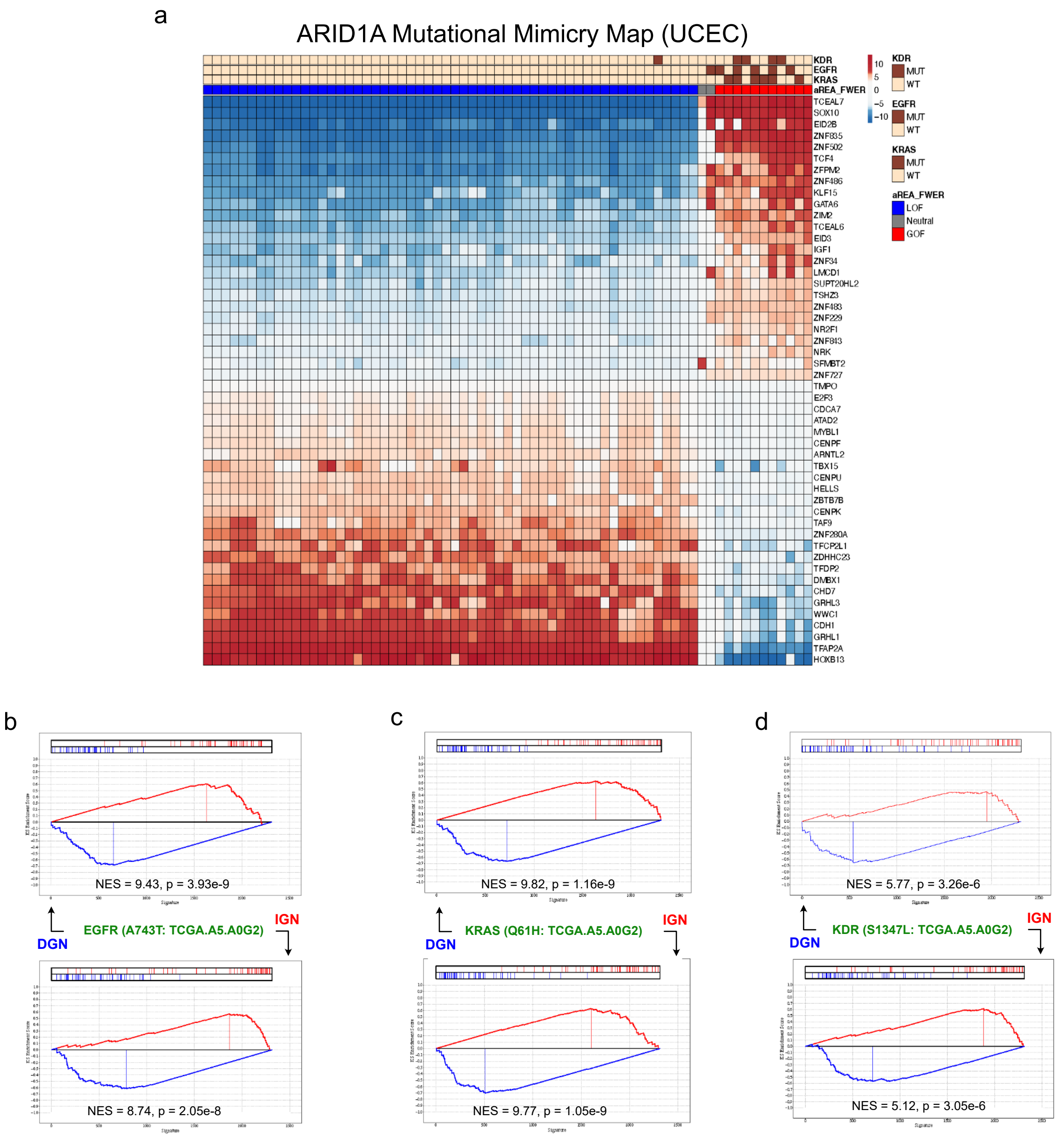
